# Direct interaction with the BRD4 carboxyl-terminal motif (CTM) and TopBP1 is required for human papillomavirus 16 E2 association with mitotic chromatin and plasmid segregation function

**DOI:** 10.1101/2023.05.25.542291

**Authors:** Apurva T. Prabhakar, Claire D. James, Christian T. Fontan, Raymonde Otoa, Xu Wang, Molly L. Bristol, Ronald D. Hill, Aanchal Dubey, Shwu-Yuan Wu, Cheng-Ming Chiang, Iain M. Morgan

## Abstract

During the human papillomavirus 16 life cycle, the E2 protein binds simultaneously to the viral genome and host chromatin throughout mitosis, ensuring viral genomes reside in daughter cell nuclei following cell division. Previously, we demonstrated that CK2 phosphorylation of E2 on serine 23 promotes interaction with TopBP1, and that this interaction is required for optimum E2 mitotic chromatin association and plasmid segregation function. Others have implicated BRD4 in mediating the plasmid segregation function of E2 and we have demonstrated that there is a TopBP1-BRD4 complex in the cell. We therefore further investigated the role of the E2-BRD4 interaction in mediating E2 association with mitotic chromatin and plasmid segregation function. Using a combination of immunofluorescence and our novel plasmid segregation assay in U2OS and N/Tert-1 cells stably expressing a variety of E2 mutants, we report that direct interaction with the BRD4 carboxyl-terminal motif (CTM) and TopBP1 is required for E2 association with mitotic chromatin and plasmid segregation. We also identify a novel TopBP1 mediated interaction between E2 and the BRD4 extra-terminal (ET) domain *in vivo*. Overall, the results demonstrate that direct interaction with TopBP1 and the BRD4 CTM are required for E2 mitotic chromatin association and plasmid segregation function. Disruption of this complex offers therapeutic options for targeting segregation of viral genomes into daughter cells, potentially combatting HPV16 infections, and cancers that retain episomal genomes.

**Importance:** HPV16 is a causative agent in around 3-4% of all human cancers and currently there are no anti-viral therapeutics available for combating this disease burden. In order to identify new therapeutic targets, we must increase our understanding of the HPV16 life cycle. Previously, we demonstrated that an interaction between E2 and the cellular protein TopBP1 mediates the plasmid segregation function of E2, allowing distribution of viral genomes into daughter nuclei following cell division. Here, we demonstrate that E2 interaction with an additional host protein, BRD4, is also essential for E2 segregation function, and that BRD4 exists in a complex with TopBP1. Overall, these results enhance our understanding of a critical part of the HPV16 life cycle and presents several therapeutic targets for disruption of the viral life cycle.

## Introduction

Papillomavirus E2 functions are mediated by three structural domains: a carboxy-terminal DNA-binding domain (DBD) that forms homodimers and binds to 12bp palindromic target sequences; an amino-terminal domain that serves as an interacting platform for host proteins and the viral helicase E1; and an unstructured hinge region that links the amino and carboxyl domains (1). E2 has three critical roles during papillomavirus life cycles. First, E1 interaction with the E2 amino-terminal domain recruits the viral helicase to the origin of replication, promoting formation of di-hexameric E1 complex that regulates viral replication in association with host proteins (2). Second, E2 can regulate transcription from the viral and host genomes in order to promote the viral life cycle (3–7). Third, E2 acts as a plasmid segregation factor that locates viral genomes into daughter nuclei following cell division (8). Our understanding of the host interacting proteins that mediate E2 functions remains incomplete and this report will focus on enhancing our understanding of the plasmid segregation function of HPV16 E2.

During mitosis, the nuclear envelope breaks down and reforms following chromosome segregation, generating nuclei in daughter cells. The viral E2 protein acts as a “bridge” during mitosis; the carboxyl terminal domain binds the viral genome while the amino-terminal interaction domain binds mitotic chromatin. The interaction with mitotic chromatin is via the E2 amino-terminal domain interacting with a host mitotic protein(s). An interaction between bovine papillomavirus 1 E2 (BPV1 E2) and BRD4 promotes the interaction between BPV1 E2 and mitotic chromatin (9–14). The interaction between all E2 proteins tested and BRD4 mediates the transcriptional regulation properties of E2 (15–19). In addition, BRD4 interaction is implicated in regulating the replication properties of E2 (20–23), and BRD4 interaction regulates the stability of E2 (24–28). Therefore, BRD4 plays a critical role during papillomavirus life cycles, partially due to a direct interaction with E2 proteins. However, the role of BRD4 in regulating the mitotic interaction and plasmid segregation function of HPV16 E2 protein is less clear. Some reports indicated that BRD4 was not the mitotic chromatin receptor for E2 (29, 30). However, other reports suggested a direct role for BRD4 in mediating HPV16 E2 association with mitotic chromatin (E2 will mean HPV16 E2 from now on unless stated otherwise) (31, 32).

Recently, we demonstrated that complexing with TopBP1 is critical for E2 interaction with mitotic chromatin and plasmid segregation (33). We initially discovered the E2-TopBP1 interaction via yeast two-hybrid screen (34, 35). Phosphorylation of E2 on serine 23 by CK2 promotes E2 interaction with TopBP1 and interaction of E2 with mitotic chromatin (36). Interaction with TopBP1 can also regulate the DNA replication properties of E2, similarly to BRD4 (20, 23, 35, 37, 38). TopBP1 is highly active during mitosis as it is involved in multiple nucleic acid processes including DNA replication and repair (39–45). Previously we demonstrated that there is a TopBP1-BRD4 complex in human cells (36). Given the interactions between E2 and TopBP1 and BRD4, the fact that TopBP1 and BRD4 exist in the same cellular complex, and previous studies implicating BRD4 in E2 interaction with mitotic chromatin, we investigated the role of BRD4 in mediating mitotic chromatin association and the plasmid segregation function of E2.

Here we demonstrate that in U2OS cells, E2 must interact with both TopBP1 and the BRD4 CTM to associate efficiently with mitotic chromatin, and in U2OS and N/Tert-1 cells these interactions are required for E2 plasmid segregation function. The amino-terminal domain of E2 binds to the carboxyl-terminal motif (CTM) of BRD4 (46), and also to TopBP1 (33, 36). Previously, we demonstrated that interaction with TopBP1 is required for E2 stabilization during mitosis and here we demonstrate that interaction with the BRD4 CTM is not required for this mitotic stabilization. We demonstrate that the interaction between BRD4 and TopBP1 is mediated by the BRD4 ET domain, and that E2 can also complex with the BRD4 ET domain via binding to TopBP1. The BRCT6 domain of TopBP1 directly interacts with the ET domain of BRD4 (47). Overall, the results demonstrate that E2 must bind directly with TopBP1 and the CTM domain of BRD4 to associate with mitotic chromatin and segregate plasmids.

## Results

### A complex interaction between E2 and TopBP1-BRD4

To investigate the roles of TopBP1 and BRD4 interaction with E2 in mitotic chromatin association and plasmid segregation function, we generated several cell lines expressing wild type and mutant E2 proteins. U2OS cells are ideal for investigating mitotic events, while N/Tert-1 cells are foreskin keratinocytes immortalized by hTERT and therefore more biologically relevant to the viral life cycle. In both cell lines we generated the following stable pools of cells: Vec, representing pcDNA vector control cells (G418 selected, as all lines were); E2-WT expresses wild-type E2; E2-S23A cells express an E2 mutant with serine 23 mutated to alanine and is compromised in TopBP1 binding (36); E2-R37A cells express an E2 mutant with arginine 37 mutated to an alanine and is proposed to be compromised in BRD4 binding (46); E2-S23A+R37A cells express a double mutant that is proposed to bind neither TopBP1 nor BRD4. Figures 1A and 1C demonstrates expression of the E2 proteins, TopBP1, BRD4-L (long form) and BRD4-S (short form) in N/Tert-1 and U2OS cells, respectively. BRD4-S is generated via differential splicing and can oppose BRD4-L functions and promote oncogenesis (48, 49). The differential splicing results in loss of the CTM, but retention of the extra-terminal (ET) domain in BRD4-S. Figures 1B and 1D represent an immunoprecipitation with E2 followed by immunoblotting for the indicated proteins. E2-WT (lane 3 in 1B and 1D) pulls down TopBP1 and both forms of BRD4. E2-S23A (lane 4) fails to bind TopBP1 as previously reported, but retains interaction with both forms of BRD4. E2-R37A (lane 5) retained interaction with TopBP1 but could also interact with both forms of BRD4. E2-S23A+R37A lost interaction with TopBP1 and both BRD4 forms (lane 6). This latter result suggested that TopBP1 might mediate the interaction between E2-R37A and both forms of BRD4 as this interaction is lost when E2-R37A fails to interact with TopBP1 (i.e., the E2-S23A+R37A mutant).

**Figure 1.**
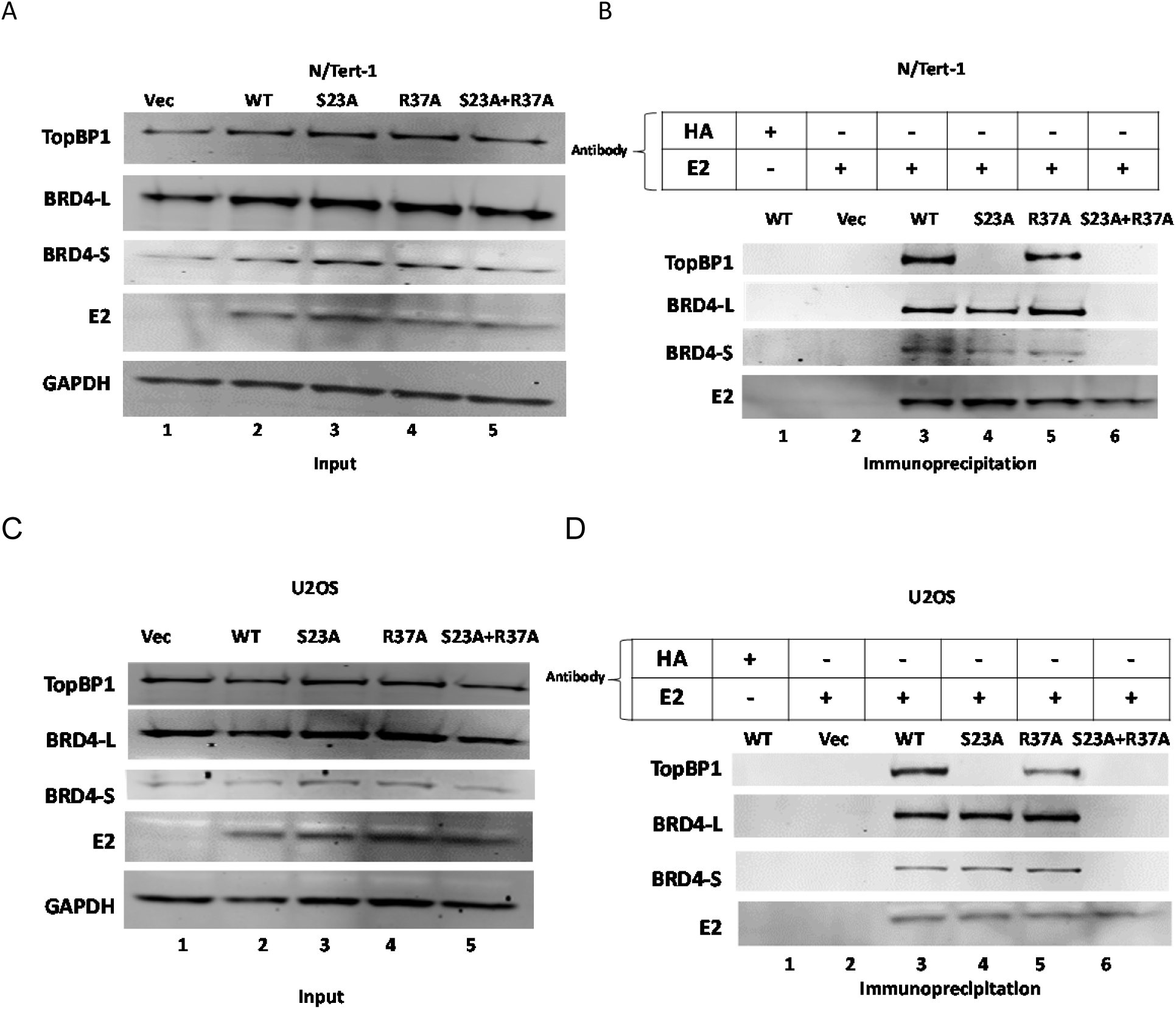
E2-R37A retains wild type interaction with BRD4 in co-immunoprecipitation. A. and C. N/Tert-1 and U2OS cell lines stably expressing the indicated E2 proteins were generated. B. and D. Immunoprecipitation of E2 brings down the E2 protein and the associated BRD4 and TopBP1 proteins with wild type and mutant E2 proteins.

To test whether E2-R37A retains BRD4 interaction via TopBP1, we knocked down TopBP1 expression using siRNA (Figure 2). Figure 2A demonstrates that the knockdown of TopBP1 (right-hand panel) did not have a significant effect on E2 expression when compared with the siRNA control samples (left-hand panels). Immunoprecipitation with an antibody that recognized only the long form of BRD4 in the siRNA control samples pulled down BRD4, along with TopBP1 and E2-WT, E2-S23A and E2-R37A, but not E2-S23A+R37A (lanes 3, 4, 5 and 6, respectively; left-hand panel). This agrees with the data in Figure 1, where E2 co-IPs demonstrated that E2-R37A could bind to BRD4, while E2-S23A+R37A could not. However, when TopBP1 is knocked down (Figure 2B, right hand panel) E2-R37A does not co-immunoprecipitate with BRD4 (lane 5), supporting a model in which E2-R37A binds to BRD4 via interaction with TopBP1. To further investigate this interaction, we carried out immunoprecipitations with an E2 antibody (Figure 2C). Again, E2-WT, E2-S23A and E2-R37A could all bind BRD4-L and -S forms, while E2-S23A+R37A could not (lanes 3, 4, 5 and 6 respectively; left-hand panel). When TopBP1 is knocked down (right-hand panel), E2-WT and E2-S23A retained interaction with BRD4-L, which is lost in E2-R37A and E2-S23A+R37A (lanes 3, 4, 5 and 6, respectively). When TopBP1 is knocked down, none of the E2 proteins interact with BRD4-S. This demonstrates that the interaction between E2 and BRD4-S only occurs due to co-binding of BRD4-S and E2 to TopBP1. TopBP1 BRCT 6 and BRD4 ET domains are the interacting domains for these two proteins, and the ET domain is retained on BRD4-S (47). The conclusion from these results is that in N/Tert-1 cells E2 only interacts directly and primarily with one domain of BRD4, the CTM, which is only on BRD4-L. We repeated these experiments in U2OS cells, with very similar results demonstrating that E2 interacts primarily with the CTM of BRD4 in these cells also (Figure 2D-2F). We repeated these experiments with an additional TopBP1 siRNA with identical results (Figure S1).

**Figure 2.**
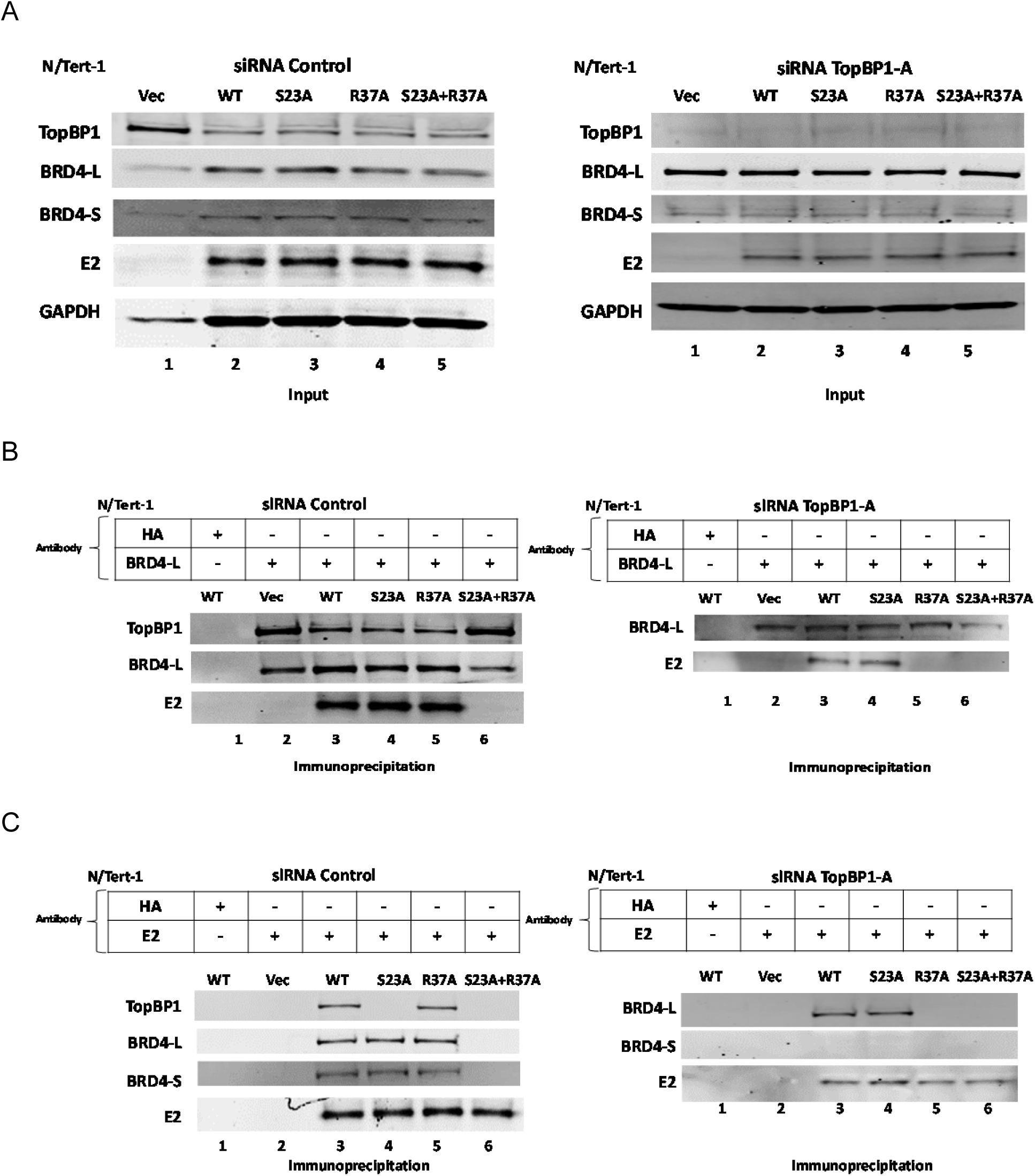
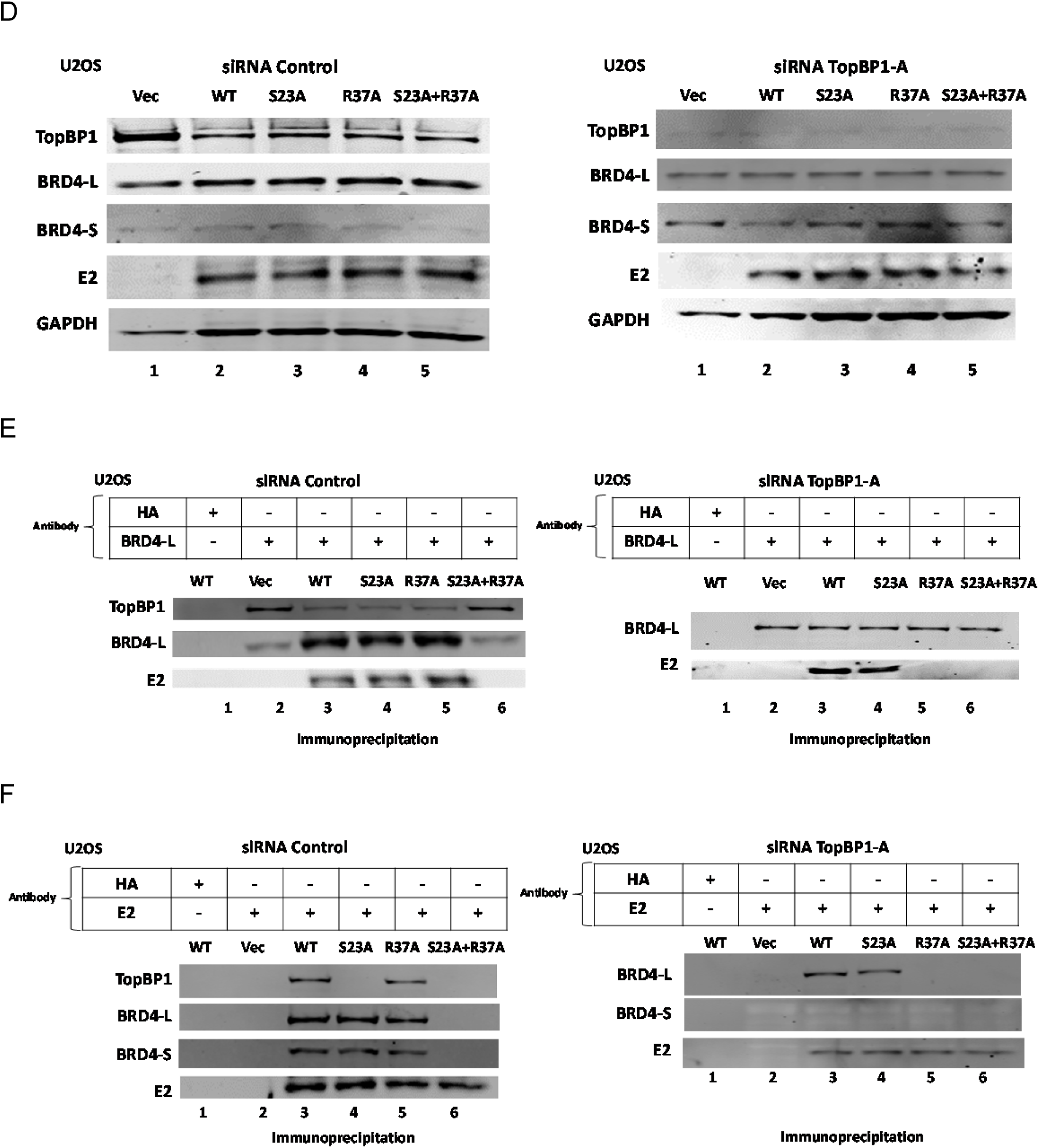
E2-R37A interacts with BRD4 via TopBP1. A. and D. Western blots demonstrating input protein levels for the immunoprecipitation experiments for N/Tert-1 and U2OS cells, respectively. Protein levels are shown in the presence of wild type (left hand panel) and siRNA knock-down of TopBP1 (right hand panel). B. and E. Cell extracts were immunoprecipitated with a BRD4 antibody that recognizes only the full length BRD4-L and the presence of the indicated proteins determined using western blots. C. and F. Cell extracts were immunoprecipitated with an E2 antibody and the presence of the indicated proteins determined using western blots.

To confirm that the ET domain of BRD4 mediated the interaction with TopBP1 and E2-R37A, we transfected FLAG-BRD4 WT (long-form) and FLAG-BRD4 ΔET into N/Tert1 and U2OS cells and investigated the ability of these FLAG-tagged BRD4 proteins to interact with the E2 proteins (Figure 3). In N/Tert-1 (Figure 3A) and U2OS (Figure 3C) the expression of the FLAG tagged BRD4 proteins had no effect on E2 or TopBP1 levels when compared with the FLAG control samples (expressing an “empty” FLAG tag). FLAG-BRD4 WT pulled down E2 WT, E2-S23A, E2-R37A but not E2-S23A+R37A in N/Tert-1 (Figure 3B) and U2OS (Figure 3D) cells (lanes 5, 8, 14 and 17, respectively). It also pulled down TopBP1 in all samples, including Vec control (lane 2). In both cell types FLAG-BRD4 ΔET pulled down E2-WT and E2-S23A (lanes 6 and 9) but could not pull down E2-R37A nor E2-S23A+R37A (lanes 15 and 18). FLAG-BRD4 ΔET could not pull down TopBP1 in Vec control cells (lane 3) nor in E2-R37A or E2-S23A+R37A cells (lanes 15 and 18), but could in E2-WT and E2-S23A cells (lanes 6 and 9). FLAG-BRD4 ΔET pulls down TopBP1 in both E2-WT and E2-S23A because both of these proteins will bind to endogenous BRD4 via the CTM which can then pull down endogenous TopBP1. The E2-R37A and E2-S23A+R37A fail to do this, as they cannot bind the BRD4 CTM domain. These results confirm that the ET domain of BRD4 is required for interaction with TopBP1, and that E2-R37A can complex with BRD4 via ET interaction with TopBP1 as E2-R37A can bind to TopBP1.

**Figure 3.**
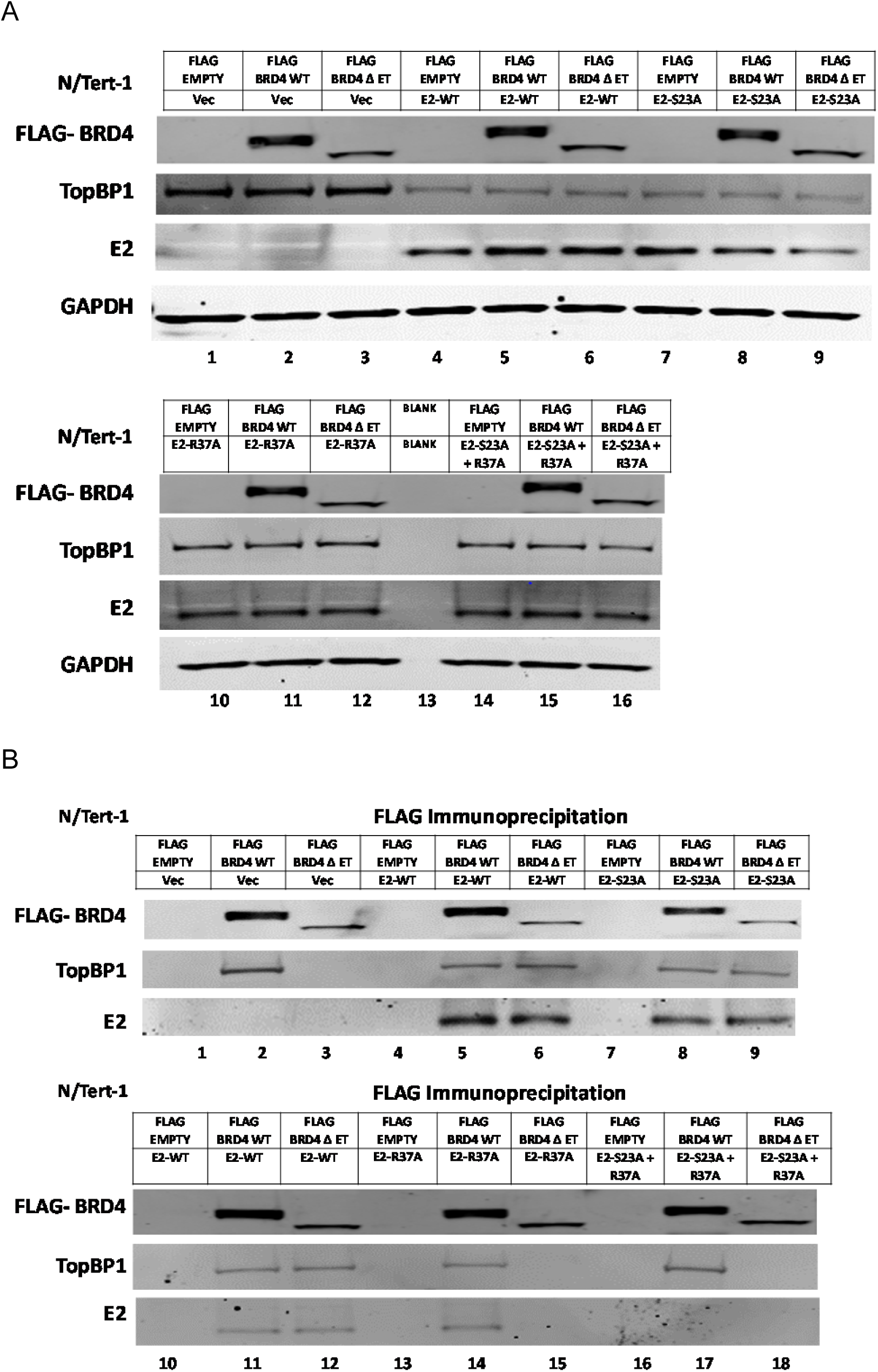
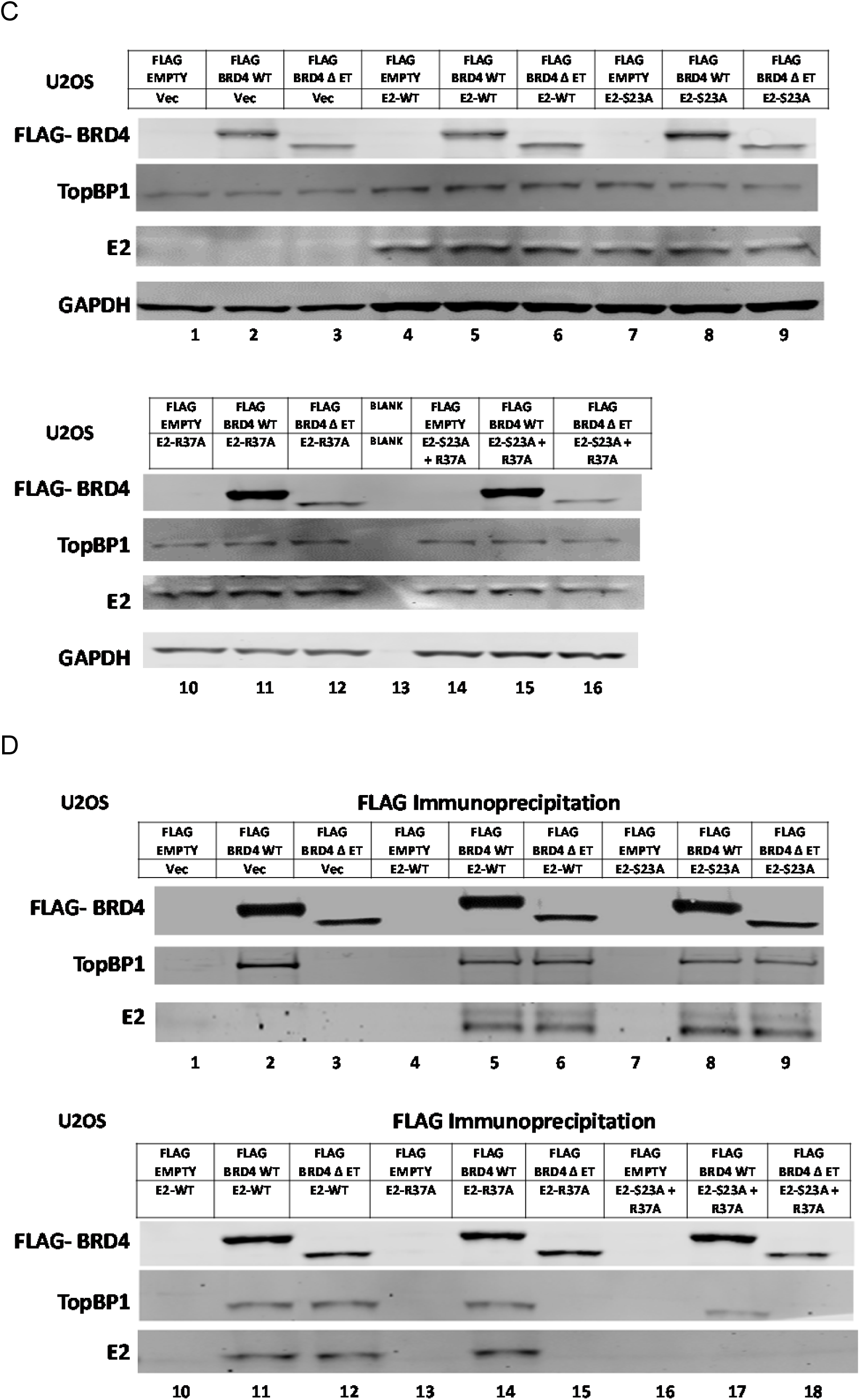
Interaction of FLAG tagged wild type (WT) BRD4 and FLAG tagged BRD4 lacking the ET domain (ΔET) with E2 proteins. A. and C. Input levels of FLAG-WT BRD4 and FLAG-ΔET BRD4 along with TopBP1 and E2 in N/Tert-1 and U2OS cells, respectively. The top panels indicate the transfected FLAG vector and the E2 protein line they were transfected into. The FLAG tagged BRD4 proteins were transiently expressed, cell extracts were prepared 48 hours following transfection. B. and D. Cell extracts were immunoprecipitated using a FLAG antibody and the indicated proteins detected by western blotting in N/Tert-1 and U2OS cells, respectively.

### A model to explain the interaction between E2 and the TopBP1-BRD4 complex

Figure 1 demonstrates that E2-R37A binds to BRD4, Figure 2 demonstrates that this interaction occurs via TopBP1 as an intermediate, and Figure 3 demonstrates that the BRD4 ET domain is required for interaction with TopBP1, and therefore for interaction with E2-R37A. The question that arises from these results is why does E2-S23A not complex with TopBP1 as it can bind to BRD4, and BRD4 can bind to TopBP1 (Figures 1 and 2)? We propose the following model (Figure 4). In Figure 4A, E2-WT interacts with TopBP1 and BRD4 in a way that blocks the TopBP1-BRD4 interaction when E2 is in the complex. In Figure 4B, E2-S23A cannot complex with TopBP1 but retains the interaction with BRD4. However, when E2-S23A is in complex with BRD4 it is unable to bind simultaneously to TopBP1 BRCT6. In this model, E2-S23A interaction with the BRD4 CTM domain alters the structure of, or masks, the BRD4 ET domain therefore blocking interaction with TopBP1. In Figure 4C, E2-R37A retains interaction with TopBP1 but has no contact directly with BRD4 (as we have demonstrated in Figures 2 and 3). In this model, the BRD4 ET domain retains the ability to interact with TopBP1 BRCT6. Therefore, following E2 or BRD4 immunoprecipitation, E2-R37A can still pull down BRD4 via TopBP1 interaction (as shown in Figures 1 and 2). In Figure 4D, the E2-S23A+R37A mutant has lost the ability to interact with either TopBP1 or BRD4. In all situations, there is detectable interaction between TopBP1 and BRD4 as there will be TopBP1 and BRD4 molecules not interacting with E2.

**Figure 4.**
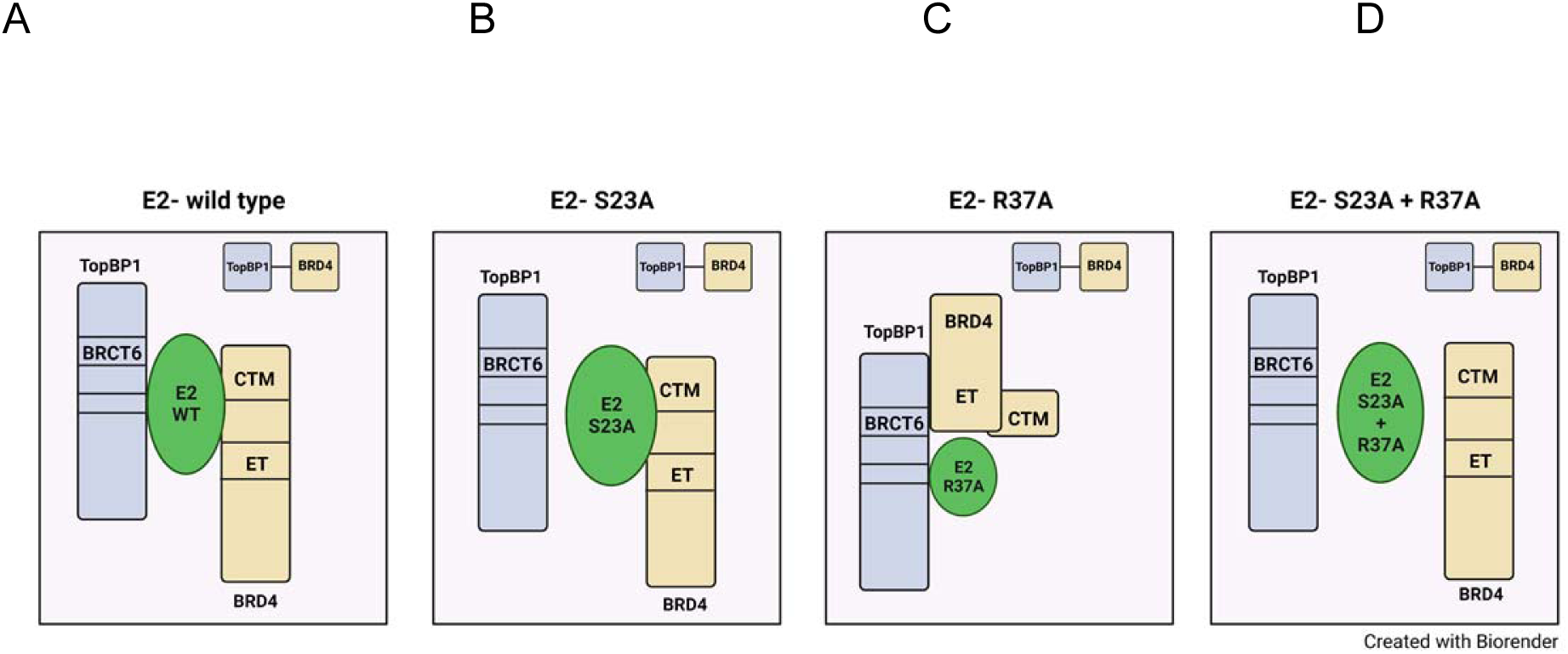
A model to explain the results presented in Figures 1-3. Please see text for details.

### Interaction with the BRD4 CTM is not required for E2 stabilization during mitosis

The interaction between E2 and TopBP1 is required for E2 stabilization during mitosis and plasmid segregation function (33, 36). To determine whether E2 interaction with the CTM of BRD4 is required for stabilization at mitosis, E2-WT, E2-S23A, E2-R37A and E2-S23A+R37A cells were synchronized into mitosis using double thymidine blocking followed by release (Figure 5). This was done in N/Tert-1 (Figure 5A and 5B) and U2OS (Figure 5C and 5D) cells. Elevated expression of cyclin B1 demonstrated mitotic enrichment. Lane 6 in Figure 5A and 5C demonstrates that E2-WT and TopBP1 protein levels increase during mitosis while lane 9 demonstrates that neither TopBP1 nor E2-S23A is stabilized during mitosis. We have reported this previously (33, 36). In both N/Tert-1 and U2OS cells TopBP1 and E2-R37A proteins levels increase in mitosis (lane 12 in Figure 5A and 5C). With E2-S23A+R37A, neither TopBP1 nor E2 protein levels increase during mitosis. These results were repeated and quantitated (Figures 5B and 5D). The results demonstrate that E2 interaction with the BRD4 CTM is not required for mitotic stabilization. None of the changes in TopBP1 or E2 protein levels are due to changes in RNA levels (33).

**Figure 5.**
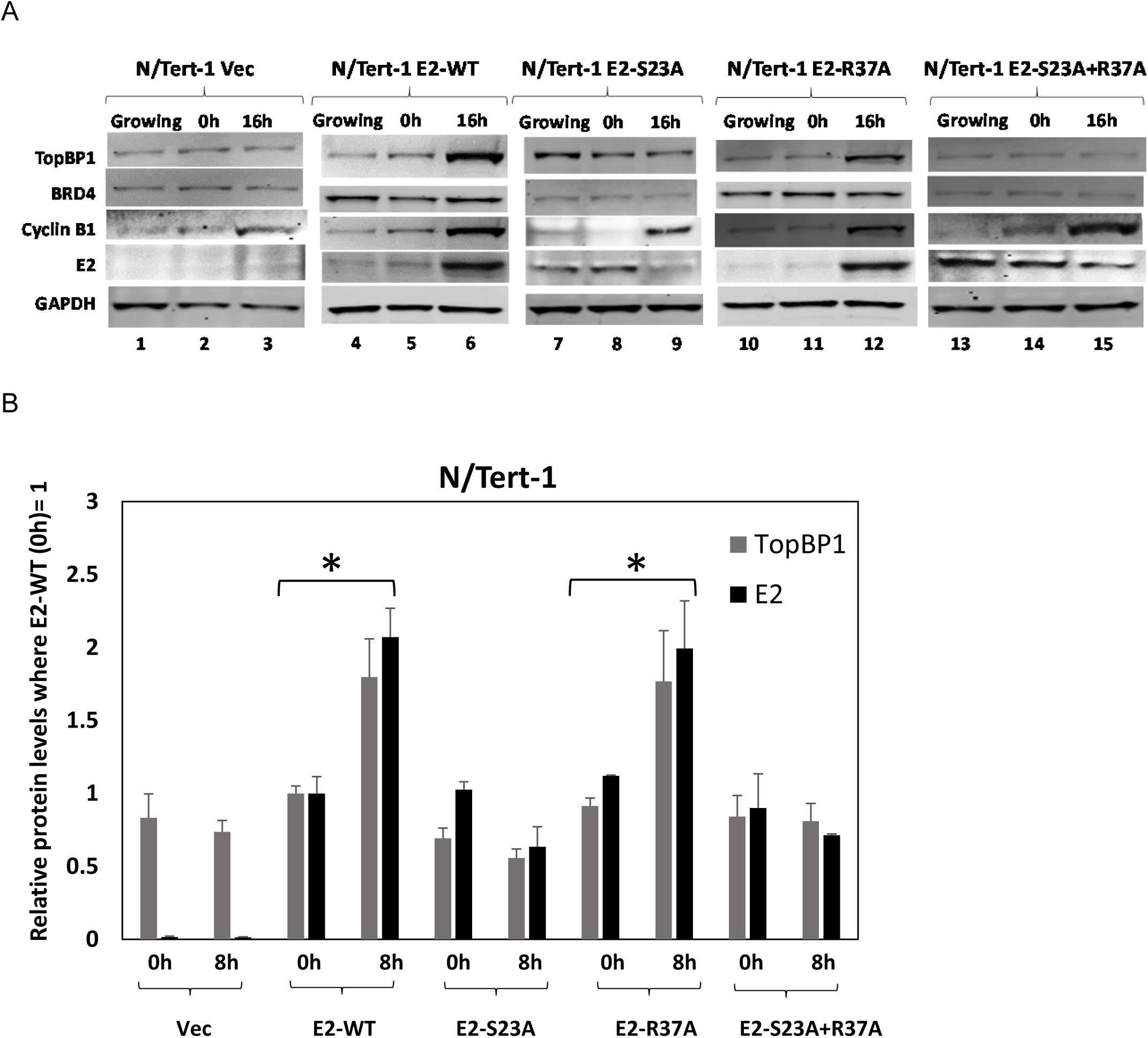
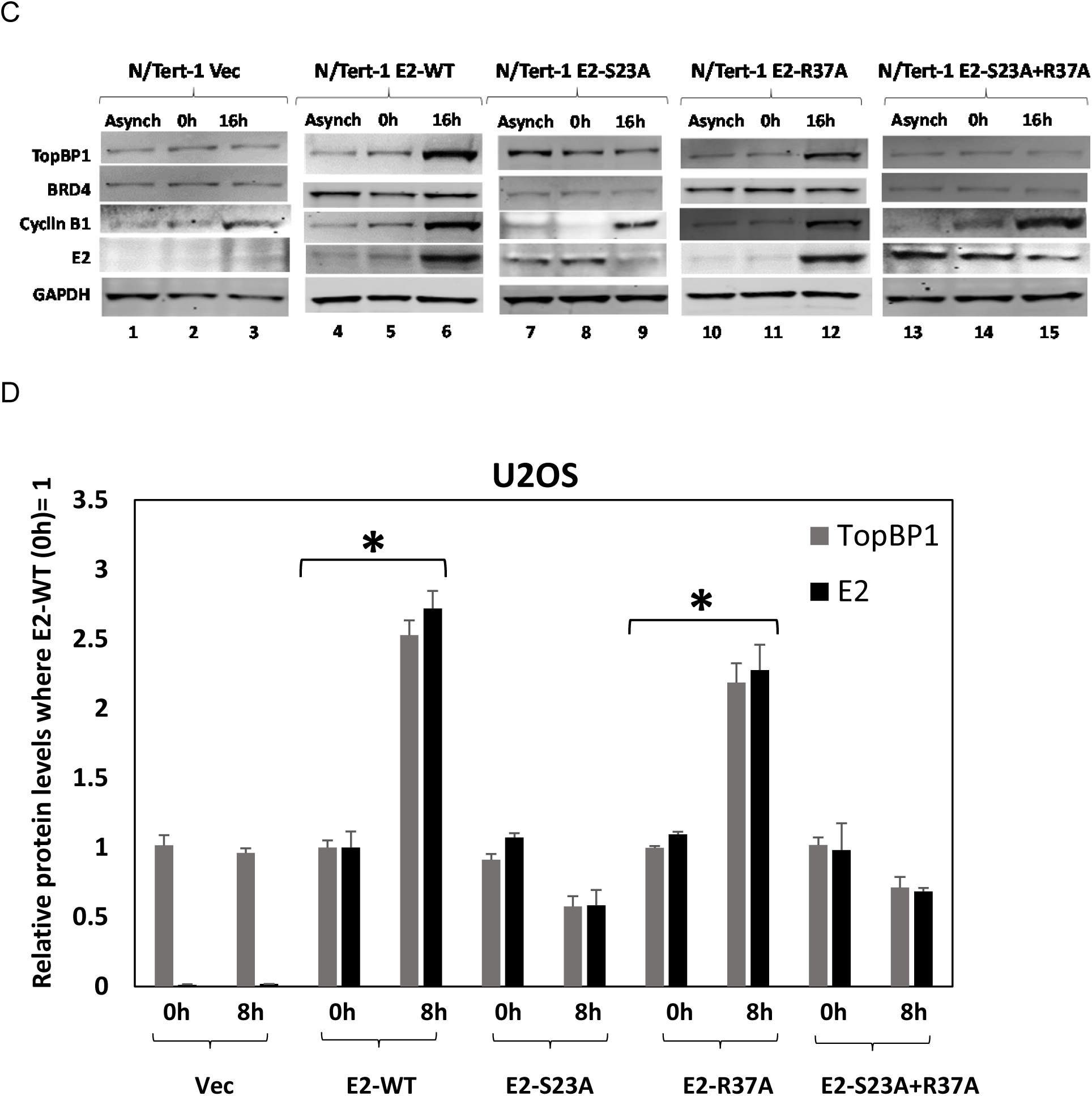
E2-R37A protein levels increase during mitosis. A. and C. The levels of the indicated proteins in N/Tert-1 and U2OS, respectively, were determined in asynchronous cells (Asynch), cells arrested by double thymidine blocking (0h) and cells released into mitosis (16 hours for N/Tert-1 cells and 8 hours for U2OS cells). Cyclin B levels confirm the enrichment of mitotic cells in the 16h and 8h samples. B. and D. The experiments were repeated and quantitated. An asterisk indicates a *P* value of less than 0.05 for the difference between the cells arrested by double thymidine blocking (0h) and cells released into mitosis (16 hours for N/Tert-1 cells and 8 hours for U2OS cells).

### E2 interaction with the BRD4 CTM is required for E2 interaction with mitotic chromatin and plasmid segregation function

Figure 6 shows images from proliferating U2OS cells fixed and stained with E2 and TopBP1 antibodies (Figure 6A) or E2 and BRD4 antibodies (Figure 6B). Mitotic cells are highlighted with white arrows in both figures. During mitosis, both TopBP1 and BRD4 co-locate with mitotic chromatin irrespective of E2 expression. E2-WT also co-locates with mitotic chromatin, as we have demonstrated previously (33, 36) (second panel down in Figures 6A and 6B). E2-S23A fails to co-locate to mitotic chromatin (third panel down in Figures 6A and 6B), as does E2-R37A (fourth panel down). Finally, E2-S23A+R37A is excluded from co-localization with mitotic chromatin (bottom panel) and the phenotype is stronger than E2-S23A or E2-R37A individually where both retain some residual interaction with mitotic chromatin. The results demonstrate that interaction with both TopBP1 and the BRD4 CTM is required for E2 recruitment to mitotic chromatin. Even though E2-R37A is stabilized during mitosis (Figure 5), this is not sufficient for E2 localization to mitotic chromatin. We next wanted to investigate whether the failure of E2 to locate to mitotic chromatin correlates with a loss of plasmid segregation function.

**Figure 6.**
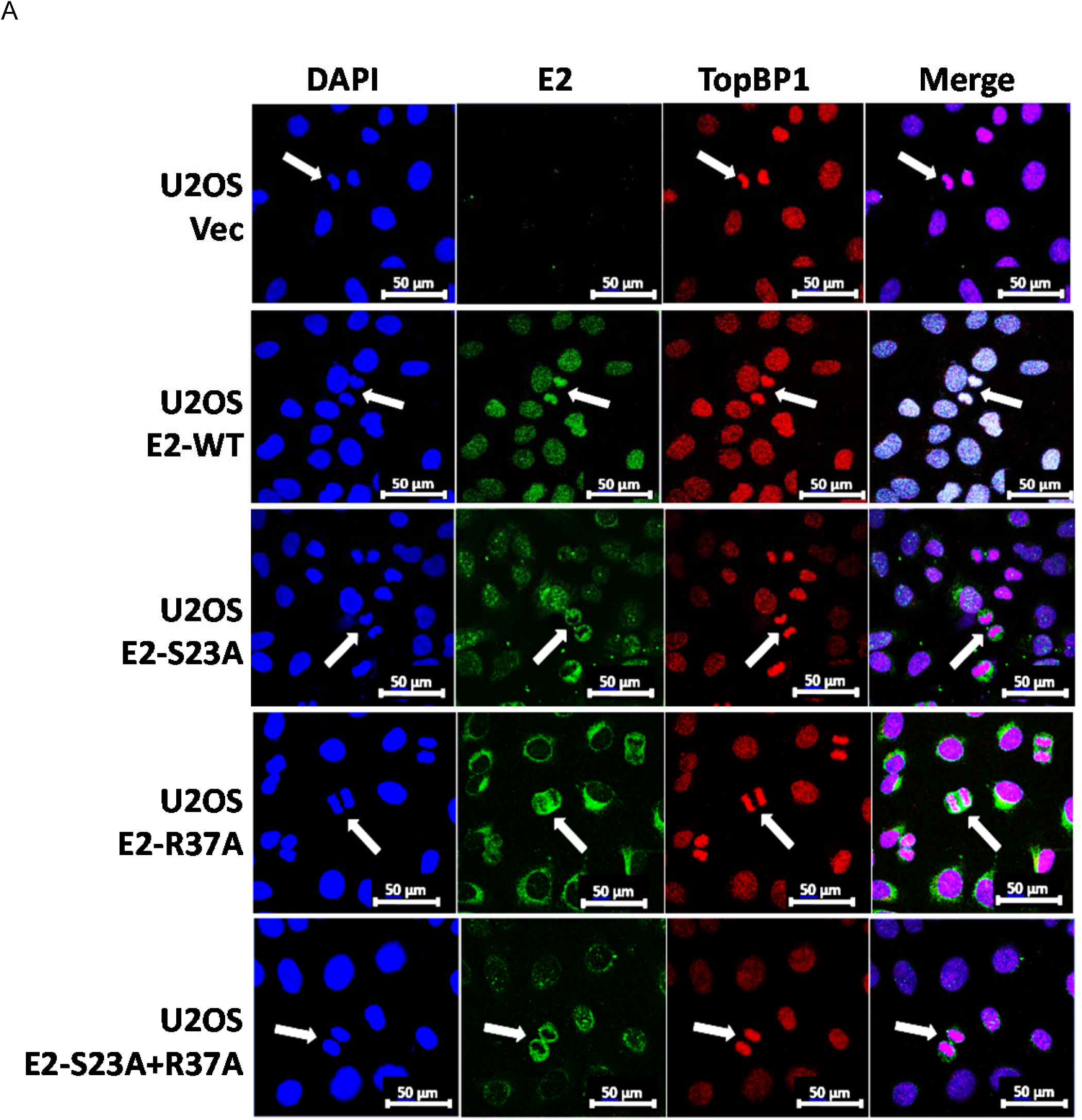
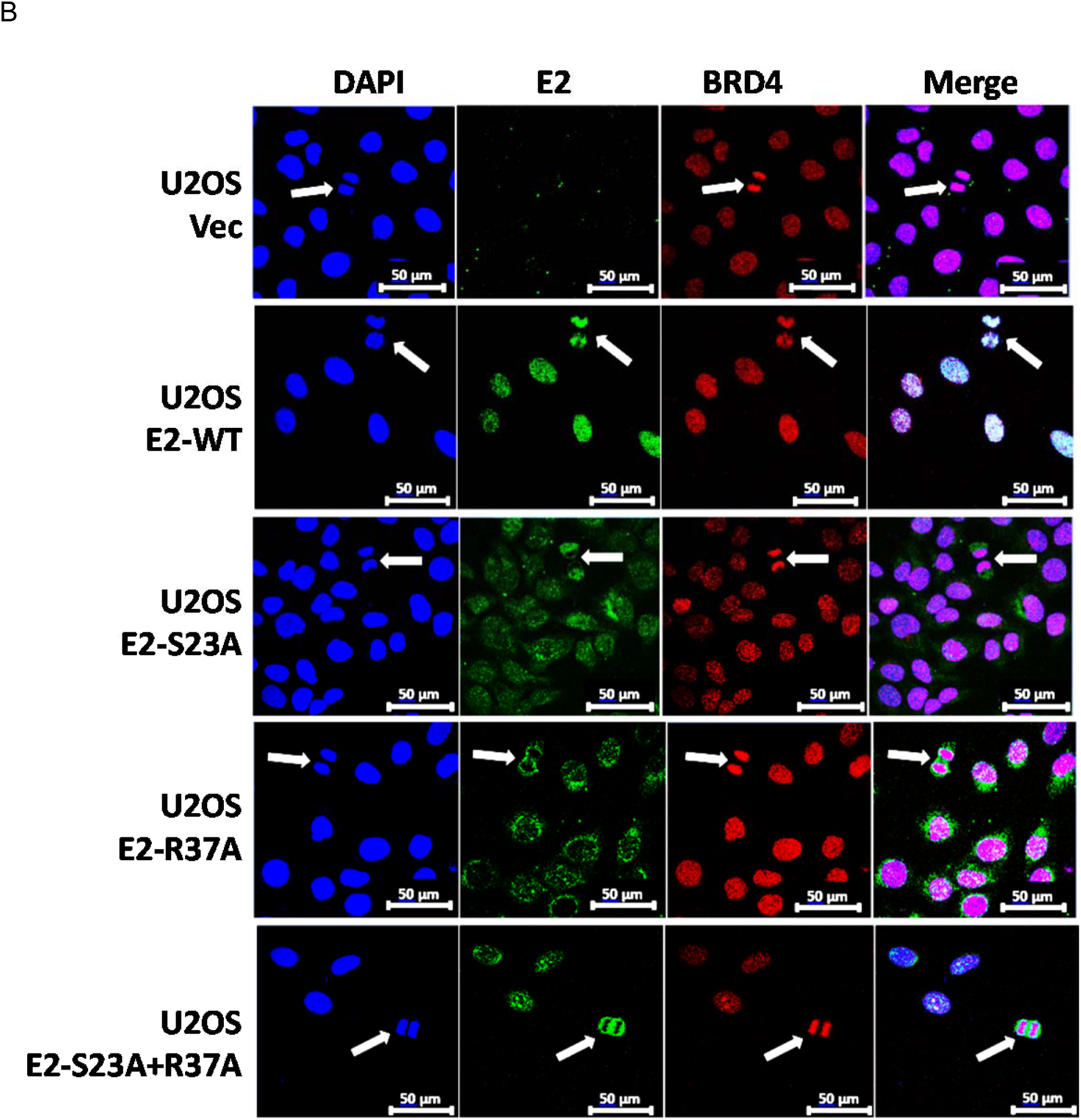
E2-R37A nor E2-S23A interact efficiently with mitotic chromatin. A. The U2OS cell lines were fixed and stained with DAPI, E2 and TopBP1. B. The U2OS cell lines were fixed and stained with DAPI, E2 and BRD4. The cell lines were asynchronously growing and mitotic cell images captured following scanning. White arrows indicate mitotic cells.

Recently, we published a report detailing a novel assay for measuring E2 plasmid segregation function (33). In this assay, plasmid DNA is labelled in a test tube and transfected into cells and the fluorescent signal generated from the label is monitored. Figure 7A represents images from the indicated U2OS cells three days following transfection with a fluorescently labelled pHPV16LCR-luc; a plasmid containing the HPV16 LCR placed upstream from the luciferase gene. The LCR contains four E2 DNA binding sites. The second panels down demonstrate that pHPV16LCR-luc co-locates with mitotic chromatin in U2OS E2-WT cells; with all of the mutant E2 protein expressing lines there is no indication of recruitment of the fluorescent plasmid to mitotic DNA. To quantitate this we scanned for the number of positively staining red cells at days 3, 6 and 9 following transfection, as we have described (33). Following days 3 and 6 the cells are split to allow for continued cell growth and mitotic events that would promote the loss of the fluorescent plasmid from the transfected cells. The experiment was repeated and the fluorescence was quantitated and graphed (Figure 7B). At day 3 following transfection over 80% of the E2-WT cells have a fluorescent signal which decreases at day 6 and 9. The fluorescent signal detected in the mutant E2 cell lines is significantly reduced compared with E2-WT. The E2-S23A+R37A mutant is the poorest at retaining the fluorescent plasmid, correlating with the complete failure of this E2 protein to co-locate to mitotic chromatin (Figure 6). Previously we have demonstrated that non-E2 containing plasmids are not located to mitotic chromatin in any of the U2OS cells, nor are they retained over an extended time period (33). In addition, we cannot co-stain these cells for E2 as the antibody permeabilization step results in loss of the fluorescent plasmid. However, there is a clear correlation between E2 interaction with mitotic chromatin (Figure 6) and plasmid retention function (Figure 7A and 7B).

**Figure 7.**
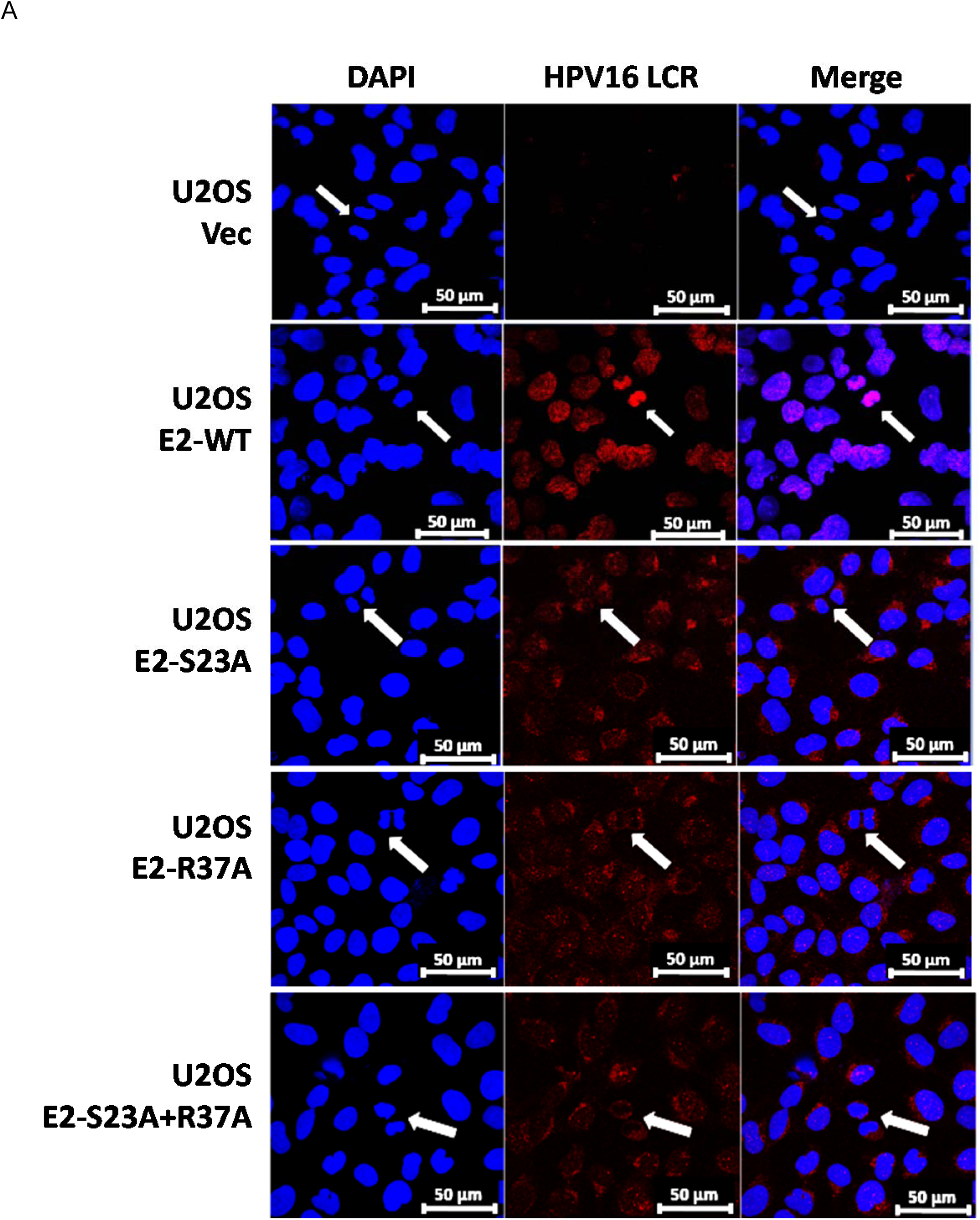
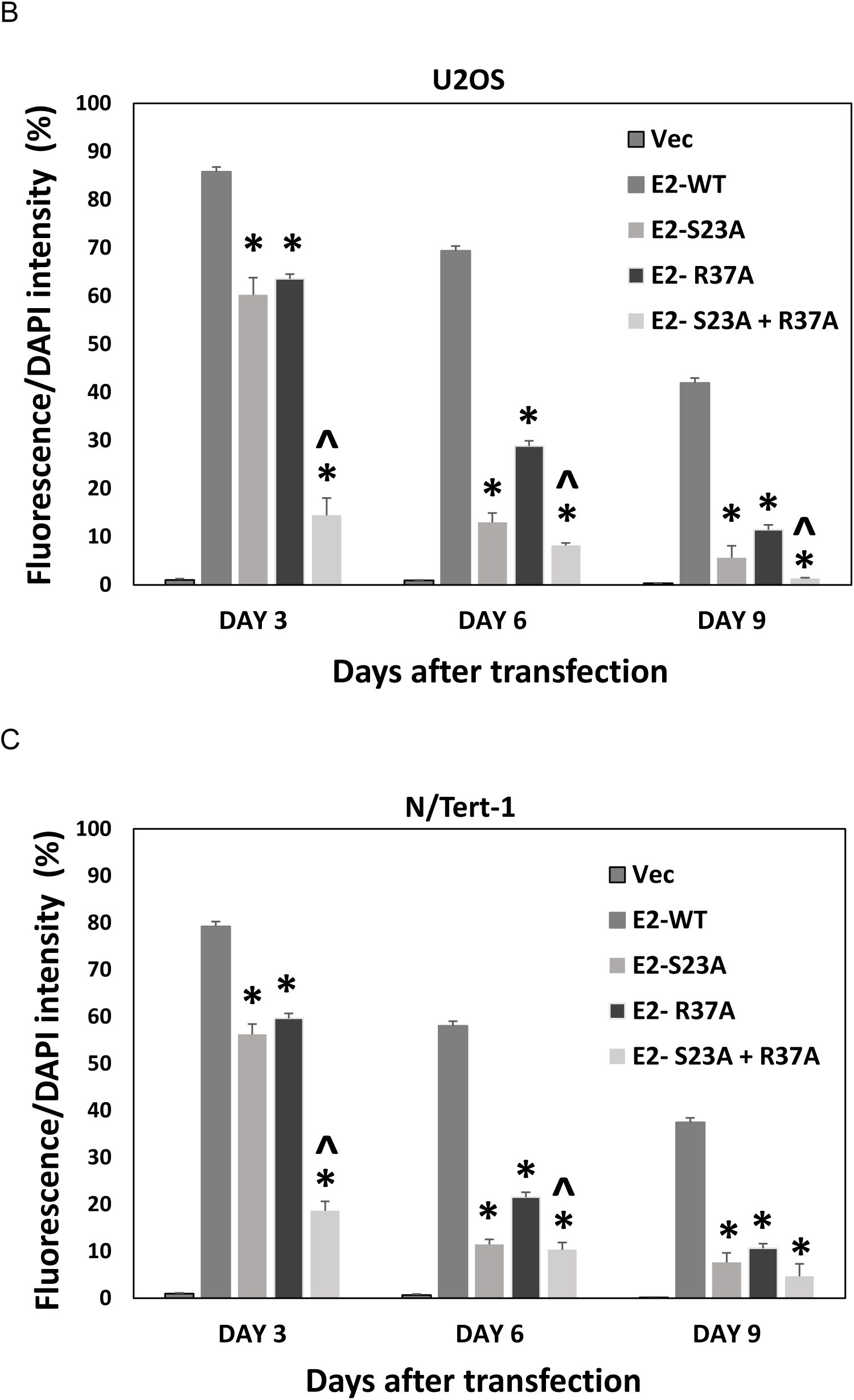
E2-R37A and E2-S23A are compromised in plasmid segregation function. A. The indicated U2OS cells were transiently transfected with a fluorescently labelled pHPV16LCR-luc and images captured 3 days following transfection. White arrows indicate mitotic cells. B. Quantitation of fluorescent plasmid retention in U2OS cells over a 9-day period following transfection. C. Quantitation of fluorescent plasmid retention in N/Tert-1 cells over a 9-day period following transfection. For both cell lines cells were trypsinized and replated on day 3 and fluorescence analyzed at day 6. This was repeated with the day 6 cells and fluorescence measured at day 9. (*) indicates a *P* value of less than 0.05 for the difference between E2-WT and other individual E2-mutant samples. (^) indicates a *P* value of less than 0.05 for the difference between E2-R37A and E2-S23A+R37A samples.

We were unable to image mitotic N/Tert-1 cells, perhaps because they become less adherent during mitosis and are lost from the coverslips during gentle washing and fixing. However, we were able to carry out our plasmid segregation assay in N/Tert-1 as this measures fluorescent signal retention (33). Therefore, we carried out our quantitative plasmid segregation assay over a 9-day period in N/Tert-1 cells and the summary of repeat experiments is shown in Figure 7C. The results in N/Tert-1 are very similar to those in U2OS cells, E2-WT is statistically better at retaining fluorescence at all time points and the E2-S23A+R37A mutant is the most compromised in retaining fluorescence. This result demonstrates that optimal E2 interaction with the TopBP1-BRD4 complex is required for E2 plasmid segregation function in human keratinocytes.

## Discussion

A breakthrough in our understanding of how papillomaviruses segregate their viral genomes during mitosis came with the discovery of an interaction between BPV1 E2 and BRD4 that tethered viral genomes to mitotic chromatin (9). Disruption of this interaction resulted in viral genome loss and phenotypic reversion of BPV1 transformed cells, further supporting a critical role for the BPV1 E2 interaction with BRD4 in mediating viral genome segregation (50). Although alpha-HPV E2 proteins retained interaction with BRD4, the role of this interaction in mediating E2 binding to mitotic chromatin and mediating plasmid segregation function remained unclear. Reports suggested that HPV16 E2 interaction with mitotic chromatin was independent of interaction with BRD4, while others indicated that interaction between HPV16 E2 and BRD4 was important for this function (30–32). These studies depended upon transient induction of E2 proteins, or transient transfection of E2 expression plasmids. We sought to resolve the role of BRD4 in mediating HPV16 E2 interaction with mitotic chromatin and segregating E2 binding site containing plasmids using cell lines stably expressing E2 proteins. The levels of E2 proteins present in our model U2OS and N/Tert-1 cells are similar to levels detected in human foreskin keratinocytes immortalized by the entire HPV16 genome (51).

Recently, we demonstrated that CK2 phosphorylation of E2 on serine 23 promotes interaction with TopBP1 *in vitro* and *in vivo* (36). Failure to interact with TopBP1 resulted in reduced interaction of E2 with mitotic chromatin, and we demonstrated that the E2-TopBP1 interaction was required for E2 plasmid segregation function (33). The fact that TopBP1 and BRD4 are in the same cellular complex, and the previous proposed role for BRD4 in mediating E2 segregation function, prompted us to investigate the role of BRD4 in this E2 function using our model systems. Figure 5 demonstrated that the interaction between E2 and the CTM domain of BRD4 (the E2-R37A mutant) is not required for stabilization during mitosis; interaction between E2 and TopBP1 is. Figure 6 demonstrates that, in U2OS cells, E2-R37A is compromised in interaction with mitotic chromatin when compared with E2-WT. E2-S23A is also compromised in mitotic chromatin interaction, while the E2-S23A+R37A mutant has lost all mitotic chromatin interaction. This figure was generated with asynchronously growing cells, there was no mitotic enrichment protocol. This is different from our previous report on E2-S23A interaction with mitotic chromatin (36), where the U2OS cells were treated to enrich for mitotic cells, and there are differences between the two sets of experiments. TopBP1 was shown to surround mitotic chromatin in distinct foci in the treated U2OS cells that did not express E2 (33) and this was similar to other reports investigating the mitotic chromatin interaction with TopBP1 following mitotic cell enrichment (40). However, in Figure 6 it is clear that TopBP1 “coats” mitotic chromatin, suggesting that the treatment to enrich for mitotic cells in previous studies promotes aberrant TopBP1 interaction with mitotic chromatin. Also, in treated U2OS cells, E2-S23A was still localized with mitotic chromatin but the staining was weaker due to a failure to stabilize during mitosis. In Figure 6, it is clear that E2-S23A has compromised mitotic chromatin interaction, again suggesting that the treatment enriching for mitotic cells alters E2 interaction with mitotic chromatin. BRD4 “coats” mitotic chromatin in a similar manner to TopBP1, and E2-R37A has reduced interaction with mitotic chromatin when compared with E2-WT (Figure 6). The E2-S23A+R37A double mutant is more compromised in mitotic chromatin interaction than either mutant by itself. A striking feature of E2-R37A staining in interphase cells was a significant reduction in nuclear detection, with a perinuclear pattern in most cells. Nuclear E2 remains, but there is more non-nuclear E2 protein. Interaction with BRD4 can regulate E2 retention in the nucleus and also protein stability (26, 28), and we have also demonstrated that chromatin attached E2 is more stable than soluble E2 (52). To our knowledge, this is the first time E2-R37A staining in stably expressing and asynchronously growing cells has been reported, and the cytoplasmic relocation of this protein could contribute to the inability of HPV16 genomes deficient of E2-BRD4 interaction to retain episomal viral genomes (18). Clearly, failure to locate to nuclei could also affect plasmid retention in our segregation assay.

To determine whether compromised interaction with mitotic chromatin resulted in compromised E2 plasmid segregation function, we used our recently developed assay (33). In this assay, an E2 binding site containing plasmid (pHPV16LCR-luc) is fluorescently labeled and transfected into the U2OS and N/Tert-1 cells, and fluorescence levels followed over a 9-day period. Figure 7A demonstrates recruitment of the fluorescent plasmid to mitotic chromatin only in the presence of E2-WT. Quantitation of fluorescence over the 9-day period demonstrates significantly greater fluorescence retention by E2-WT than that of any other cell line. E2-S23A+R37A was the poorest at retention, correlating with the poorest interaction with mitotic chromatin (Figure 6). This fluorescence retention required a plasmid containing E2 DNA binding sites, and was not due to integration of the fluorescent DNA (33).

The involvement of the E2-BRD4 interaction in plasmid segregation function further highlights the importance of CK2 in this process. We have already demonstrated that CK2 phosphorylation of E2 serine 23 is required for the E2-TopBP1 interaction (36). CK2 phosphorylation of BRD4 regulates BRD4 interaction with chromatin and hyper-phosphorylation of BRD4 by CK2 is linked to progression of triple-negative breast cancer (TNBC) (53, 54). CK2 phosphorylation of E7 is required for the maintenance of the E7 transformed phenotype (55), therefore targeting CK2 is an attractive therapeutic target for disrupting HPV16 infections and cancers.

The conclusion from these experiments is that E2 must directly contact TopBP1 and the BRD4 CTM to efficiently interact with mitotic chromatin and segregate plasmids during cell division. Figures 1-3 demonstrate that there is a complex interaction between E2 and BRD4-TopBP1. The BRD4 ET domain is required for interaction with TopBP1 (Figure 3) and this interaction allows E2-R37A to retain complex formation with BRD4 via TopBP1. siRNA knock-down of TopBP1 eliminates E2-R37A interaction with BRD4-L and -S forms. Previous studies demonstrated a direct interaction between the DNA binding region of E2 and the basic residue-enriched interaction domain (BID) and phospho-N-terminal phosphorylation sites (pNPS) of BRD4 (23). It is likely that intramolecular contacts between ET, along with its juxtaposed C-terminal phosphorylation sites (CPS), may regulate the conformation and extent of surface exposure of CK2-regulated pNPS that in turn modulates BID accessibility, thereby controlling selective contact with E2 in a context-specific and cell type-dependent manner. In N/Tert-1 and U2OS cells It is possible that, once recruited to mitotic chromatin, E2 can then interact with the BID and/or pNPS.

Future studies will focus on a deeper characterization of the E2-TopBP1-BRD4 complex. Enhancing our understanding of this complex will provide therapeutic targets for disrupting HPV plasmid segregation and potentially alleviating HPV infection and disease burden.

## Materials and methods

### Cell culture and plasmids

Stably expressing HPV16 wild-type E2 (E2-WT), mutants E2-S23A (E2 with serine 23 mutated to alanine, abrogating interaction with TopBP1), E2-R37A (E2 with arginine 37 mutated to alanine), double mutant E2-S23A + R37A (E2 with serine 23 mutated to alanine and arginine 37 mutated to alanine) and an empty vector plasmid control, pCDNA 3.0 were generated as previously described (33, 36). N/Tert-1 cells were cultured in keratinocyte serum-free medium (K-SFM) (Invitrogen; catalog no. 37010022) supplemented with bovine pituitary extract, EGF (Invitrogen), 0.3 mM calcium chloride (Sigma; 21115) and 150 μg/ml G418 (Thermo Fisher Scientific) cultured at 37 °C in a 5% CO_2_ / 95% air atmosphere. U2OS cells were passaged in Dulbeccos Modified Eagle Medium (DMEM) supplemented with 10% FBS and 1.5 mg/mL G418 sulfate (33, 36). pCDNA5-FLAG-BRD4-WT and pFLAG-CMV2-BRD4 ΔET plasmids were purchased from Addgene. pCDNA-Flag vector control plasmid was a kind gift from Dr. Yue Sun, Philips Institute for Oral Health Research. The Flag plasmids were transiently transfected in to N/Tert-1 cells using Lipofectamine™ RNAiMAX transfection (Invitrogen, catalog no. 13778-100) protocol and transfected in to U2OS cells using the calcium phosphate transfection protocol.

### Immunoblotting

Indicated cells were trypsinized, washed with 1X PBS and resuspended in 5X packed cell volume of mRIPA(400) buffer (50Mm Tris.HCl Ph 7.5, 400mM NaCl. 1%NP-40, 0.25% sodium deoxycholate, 1mM EDTA and protease inhibitors: 0.5 µM PMSF, 50 µg/ml Leupectin, 50 µg/ml Aprotinin, 1 µg/ml Pepstatin A and 500X dilution of phosphatase inhibitor cocktail 2 (Sigma)). Cell-buffer suspension was incubated for 30 min on ice with occasional agitation and then centrifuged for 15 min at 14,000 rcf at 4 °C. Protein concentration was determined using the Bio-Rad protein estimation assay according to manufacturer’s instructions. 100 μg protein samples were heated at 95 °C in 4x Laemmli sample buffer (Bio-Rad) for 5 min Samples were run down a Novex 4–12% Tris-glycine gel (Invitrogen) and transferred onto a nitrocellulose membrane (Bio-Rad) at 30V overnight using the wet-blot transfer method. Membranes were blocked with Odyssey (PBS) blocking buffer (diluted 1:1 with 1X PBS) at room temperature for 1h and probed with indicated 1° antibody diluted in Odyssey blocking buffer. Membranes were washed with PBS supplemented with 0.1% Tween (PBS-Tween) and probed with the indicated Odyssey secondary antibody (goat anti-mouse IRdye 800CW or goat anti-rabbit IRdye 680CW) diluted in Odyssey blocking buffer at 1:10,000. Membranes were washed three times with PBS-Tween and an additional wash with 1X PBS. Infrared imaging of the blot was performed using the Odyssey CLx Li-Cor imaging system. Immunoblots were quantified using ImageJ utilizing GAPDH as internal loading control. The following primary antibodies were used for immunoblotting in this study: monoclonal B9 1:500 (56), anti-TopBP1 1:1,000 (Bethyl, catalog no. A300-111A), anti-BRD4 1:1,000 (Bethyl, catalog no. A301-985A); anti-short form BRD4 (BRD4-S) 1:1000 (23); anti-FLAG epitope tag (DYKDDDDK) 1:500 (Thermo fisher, catalog no. PA1-984B); anti-Cyclin B1 (D5C10) XP 1:1,000 (Cell Signaling Technology, cat no. 4138); Glyceraldehyde-3-phosphate dehydrogenase (GAPDH) 1:10,000 (Santa Cruz; catalog no. sc-47724).

### Immunoprecipitation

Primary antibody of interest or a HA tag antibody (used as a negative control) was incubated in 250 μg of cell lysate (prepared as described above), made up to a total volume of 500 μl with lysis buffer supplemented with protease inhibitors and phosphatase inhibitor cocktail and rotated at 4°C overnight. The following day, 40 μl of prewashed protein A beads per sample (Sigma; equilibrated to lysis buffer as mentioned in the manufacturer’s protocol) was added to the lysate-antibody mixture and rotated for another 4 h at 4°C. The samples were gently washed with 500 μl lysis buffer by centrifugation at 1,000 rcf for 2 to 3 min. This wash was repeated four times. The bead pellet was resuspended in 4X Laemmli sample buffer (Bio-Rad), heat denatured, and centrifuged at 1,000 rcf for 2 to 3 min. Proteins were separated using a sodium dodecyl sulfate-polyacrylamide gel electrophoresis (SDS-PAGE) system and transferred onto a nitrocellulose membrane before probing for the presence of E2, TopBP1, BRD4-L or Flag tagged BRD4 proteins, as per the Western blotting protocol described above,

### Small interfering RNA (siRNA)

N/Tert-1 cells or U2OS cells were plated on a 100-mm plates. The next day, cells were transfected with 10 µM of the siRNA mentioned below. 10 µM of MISSION^®^ siRNA Universal Negative Control (Sigma-Aldrich; catalog no. SIC001) was used as a “non-targeting” control in our experiments. Lipofectamine™ RNAiMAX transfection (Invitrogen, catalog no. 13778-100) protocol was used in the siRNA knockdown. 48 hours post transfection, the cells were harvested, and the knockdown was confirmed by immunoblotting for the protein of interest as described above. All siRNAs were purchased from Sigma-Aldrich: siRNA TopBP1-A-CUCACCUUAUUGCAGGAGA(dTdT); siRNA TopBP1-B-GUAAAUAUCUGAAGCUGUA(dTdT).

### Cell synchronization

N/Tert-1 cells expressing stable E2-WT, E2-S23A, E2-R37A, E2-S23A+ R37A and pcDNA empty vector plasmid control were plated at 5 × 10^5^ density onto 100-mm plates in DMEM + 10% FBS medium. The cells were treated with 2 mM thymidine diluted in supplemented K-SFM media for 16 h. Cells were then washed 2X with PBS and recovered in supplemented K-SFM. After 8 h, to block the cells at G_1_/S phase, a second dose of 2 mM thymidine was added and the cells were incubated for 17 h. The cells were then washed twice with PBS and recovered as before at the following time points. For N/Tert-1, cells were harvested at 0 h (G_1_/S phase) and 16 h (M1 phase). The above procedure was repeated in U2OS cells which were plated at a density of 3 × 10^5^ on a 100-mm plates in DMEM media and the double thymidine blocked U2OS cells were harvested at 0 h (G_1_/S phase) and 8 h (M1 phase). The cell lysates were prepared using the harvested cells at the time points mentioned, and immunoblotting was carried out. Cyclin B1 antibody was used to confirm mitotic enrichment. The blots were quantitated using ImageJ.

### Fluorescently tagged plasmid transfection and imaging

Label IT Tracker (Mirus Bio, cat no. MIR7025) protocol was used to covalently attach a fluorescein-492 tag to pHPV16-LCR-luc (long control region containing transcriptional control elements, the origin or viral replication, and E2 DNA-binding sites) and to pSV40-luc (pGL3-Control from Promega, containing the SV40 promoter and enhancer regions but no E2 DNA-binding sites). These fluorescent plasmids were then transiently transfected into following stable cells: U2OS-Vec (vector control), U2OS-E2-WT (stably expressing wild type E2), U2OS-E2-S23A, U2OS-E2-R37A and U2OS-E2-S23A+R37A. 48h post-transfection, these transfected cells were passaged for next time point and imaged as previously described (33, 36). This was repeated using the N/Tert-1 E2-WT, E2-mutant cell lines and pCDNA vector cells.

### Immunofluorescence

U2OS cells expressing stable E2-WT, E2-S23A, E2-R37A, E2-S23A+R37A and pcDNA empty vector plasmid control were plated on acid-washed, poly-l-lysine-coated coverslips, in a six-well plate at a density of 2 × 10^5^ cells/well with 5 ml DMEM-FBS. After 48 h, the cells were washed twice with phosphate-buffered saline (PBS), fixed, and stained as previously described (33, 36). The primary antibodies used are as follows: HPV16 E2 B9 monoclonal antibody 1:500; BRD4 1:1000 (Bethyl laboratories, cat no. A700-004); TopBP1, 1:1,000 (Bethyl Laboratories, cat no. A300-111A). The cells were washed and incubated with secondary antibodies Alexa Fluor 488 goat anti-mouse (Thermo Fisher, cat no. A-11001) and Alexa Fluor 594 goat anti-rabbit (Thermo Fisher, cat no. A-11037) diluted at 1:1,000. The wash step was repeated and the coverslips were mounted on a glass slide using Vectashield mounting medium containing DAPI. Images were captured using a Zeiss LSM700 laser scanning confocal microscope and analyzed and quantitated using Zen LE software and Keyence analyzing system (BZ-X810).

### Statistical analysis

All the experiments were carried out in triplicates in each of the two different cell lines. Quantitation of cell synchronization assay and fluorescence/DAPI intensity results are represented as mean ± standard error (SE). A Student’s t test was used to determine significance.

## Acknowledgements

IMM’s work is supported by US NIH grant R01DE029471 and by the National Cancer Institute grant supporting VCU Massey Cancer Center, P30CA016059. C.-M. Chiang’s research is supported by US NIH grant 1RO1CA251698-01 and CPRIT grant RP180349.

